# In-depth characterisation of a selection of gut commensal bacteria reveals their functional capacities to metabolise dietary carbohydrates with prebiotic potential

**DOI:** 10.1101/2024.01.16.575889

**Authors:** Cassandre Bedu-Ferrari, Paul Biscarrat, Frederic Pepke, Sarah Vati, Cyril Chaudemanche, Florence Castelli, Céline Chollet, Olivier Rué, Christelle Hennequet-Antier, Philippe Langella, Claire Cherbuy

## Abstract

The microbial utilisation of dietary carbohydrates is closely linked to the pivotal role of the gut microbiome in human health. Inherent to the modulation of complex microbial communities, a prebiotic implies the selective utilisation of specific substrate, relying on the metabolic capacities of targeted microbes. In this study, we investigated the metabolic capacities of 17 commensal bacteria of the human gut microbiome toward dietary carbohydrates with prebiotic potential. First, *in vitro* experiments allowed the classification of bacterial growth and fermentation profiles in response to various carbon sources, including agave inulin, corn fiber, polydextrose and citrus pectin. The influence of phylogenetic affiliation appeared to statistically outweigh carbon sources in determining the degrees of carbohydrate utilisation. Secondly, we narrowed our focus on six commensal bacteria representative of the *Bacteroidetes* and *Firmicutes* phyla to perform an untargeted HR-LC/MS metabolomic analysis. *Bacteroides xylanisolvens*, *Bacteroides thetaiotaomicron*, *Bacteroides intestinalis*, *Subdoligranulum variabile*, *Roseburia intestinalis* and *Eubacterium rectale* exhibited distinct metabolomic profiles in response to different carbon sources. The relative abundance of bacterial metabolites was significantly influenced by dietary carbohydrates, with these effects being strain-specific and/or carbohydrate-specific. Particularly, the findings indicated an elevation in short-chain fatty acids and other metabolites, including succinate, gamma-aminobutyric acid, and nicotinic acid. These metabolites were associated with putative health benefits. Finally, an RNA-Seq transcriptomic approach provided deeper insights into the underlying mechanisms of carbohydrate metabolisation. Restricting our focus on four commensal bacteria, including *B. xylanisolvens*, *B. thetaiotaomicron, S. variabile* and *R. intestinalis*, carbon sources did significantly modulate the level of bacterial genes related to the enzymatic machinery involved in the metabolisation of dietary carbohydrates. This study provides a holistic view of the molecular strategies induced during the dynamic interplay between dietary carbohydrates with prebiotic potential and gut commensal bacteria.

## INTRODUCTION

Nutritional strategies can modulate the composition and functional activities of the gut microbiota, conferring health benefits. In this context, prebiotics define substrates that are selectively utilised by host microorganisms associated with host health, beyond *Bifidobacterium* and *Lactobacillus* species [1, 2]. Prebiotics mainly describe non-digestible carbohydrates, even though other compounds with prebiotic potential, such as polyphenols, are emerging [3–5]. As a consequence of the breakdown of carbohydrates, short-chain fatty acids (SCFA) often substantiate prebiotic health benefits [6, 7]. Inulin-type fructans, including short fructo-oligosaccharides (FOS) and longer inulin molecules, likely constitute the most accepted prebiotics in clinical interventions [8, 9] that can mediate satietogenic effects [10, 11] and regulate postprandial glycemia, insulinemia [12, 13] and normal cholesterol levels in the blood [14].

To gain insights into the interactions between prebiotics, gut microbes, and host health, efforts are made to characterise the mechanisms underlying the prebiotic effects of dietary compounds. Saccharolytic fermentation involves a multitude of carbohydrate-active enzymes (CAZymes) dedicated to the catalysis of complex material into individual carbohydrate components [7, 15]. In particular, glycoside hydrolases (GH) support the hydrolysis and/or rearrangement of glycosidic bonds, which are essential for bacterial foraging systems such as the machinery of polysaccharide utilisation loci (PUL) [16]. According to the degradation capacities of commensal bacteria, dietary carbohydrates can have disparate effects on the microbial communities. In the field of human health and nutrition, mechanistic research to clarify the selective utilisation of prebiotics is an important basis for the development of personalised nutritional and clinical strategies. In this study, *in vitro* experiments investigated the functional capacities of 17 gut commensal bacteria, beyond the popular *Bifidobacterium* and *Lactobacillus* groups, to metabolise four dietary carbohydrates with prebiotic potential, including agave inulin, corn fiber, polydextrose and citrus pectin. Furthermore, metabolomic and transcriptomic approaches allowed the characterisation of molecular mechanisms involved in the utilisation of these dietary carbohydrates by six and four selected bacteria, respectively.

## RESULTS

### Effect of dietary carbohydrates on bacterial growth parameters

The utilisation of dietary carbohydrates was investigated on 17 gut commensal bacteria, representing the four major phyla of the human gut microbiome [17]. The selected bacteria included prevalent members of a healthy human gut microbiota (Table S1) [18–20]. Among these bacteria, *Bacteroidetes* species are commonly recognised as primary degraders of polysaccharides due to their extensive enzymatic repertoires, such as *Bacteroides fragilis*, *Bacteroides intestinalis*, *Bacteroides thetaiotaomicron* and *Bacteroides xylanisolvens* (Table S1). As important acetate and/or butyrate producers, the selected *Firmicutes* species belong to two major families of the gut microbiota: *Lachnospiraceae* (*Eubacterium rectale*, *Anaerobutyricum hallii*, *Anaerostipes caccae*, *Blautia hansenii*, *Roseburia intestinalis* and *Roseburia inulinivorans*) and *Ruminococcaceae* (*Butyricicoccus pullicaecorum*, *Faecalibacterium prausnitzii, Ruminococcus bromii, Subdoligranulum variabile*). Among these bacteria, certain *Actinobacteria* (*Bifidobacterium adolescentis, Bifidobacterium catenulatum*) and *Firmicutes* species (*E. rectale, A. hallii, R. intestinalis, A. caccae, R. inulinivorans, B. pullicaecorum* and *B. hansenii*) were identified as potentially beneficial bacteria with implications for chronic diseases such as allergy, inflammation, metabolic syndrome, and immunotherapy (Table S1). In our study, we deliberately included extensively studied strains (e.g. *B. adolescentis*, *B. thetaiotaomicron, R. intestinalis*) and less well-studied strains (e.g. *B. catenulatum*, *A. caccae*, *S. variabile*) to provide a comprehensive understanding of their response to dietary carbohydrates.

In our *in vitro* experiments, we first evaluated the utilisation of dietary carbohydrates by each bacterial strains using low nutrient culture media (LNCM). The media were supplemented with agave inulin (LNCM-inulin), corn fiber (LNCM-corn-fiber), polydextrose (LNCM-polydextrose) or citrus pectin (LNCM-pectin) (Text S1 for compositional details). LNCM with glucose (LNCM-glucose) ran in parallel and served as positive control. Measurements of OD_600nm_ and pH were monitored as indicators of bacterial growth and fermentation (Fig. 1) [21]. Bacterial growth was low in non-supplemented LNCM, except for *Bacteroidetes* strains. In contrast, most bacteria in LNCM-glucose recorded substantial growth and medium acidification, except for *B. pullicaecorum, F. prausnitzii, R. bromii* and *A. muciniphila*, which did not grow under any carbon conditions. In regards to these conditions, each bacterium exhibited different capacities in utilising dietary carbohydrates. *Bacteroidetes* strains effectively grew and fermented all tested carbon sources. Notably, *B. xylanisolvens* and *B. thetaiotaomicron* displayed the highest growth and fermentations on all tested conditions, reaching OD and pH values obtained in the presence of glucose. In contrast, *Actinobacteria* and *Firmicutes* strains tested showed a more narrow range in utilising dietary carbohydrates. Thus, only inulin is able to stimulate the growth of *B. catenulatum*, at a level comparable to glucose. Strains belonging to *Lachnospiraceae* displayed higher growth and fermentations than strains belonging to *Ruminococcaceae*, except *S. variabile*. However, for *Firmicutes*, OD and pH values obtained in the presence of dietary carbohydrates remained lower than the ones obtained in the presence of glucose, except for *R. inulinivorans* and *B. hansenii* in the presence of inulin. Altogether, our data showed the impact of dietary carbohydrates included in this study varies among bacteria and is related to the phylogenetic affiliation of the strains.

**Figure 1:**
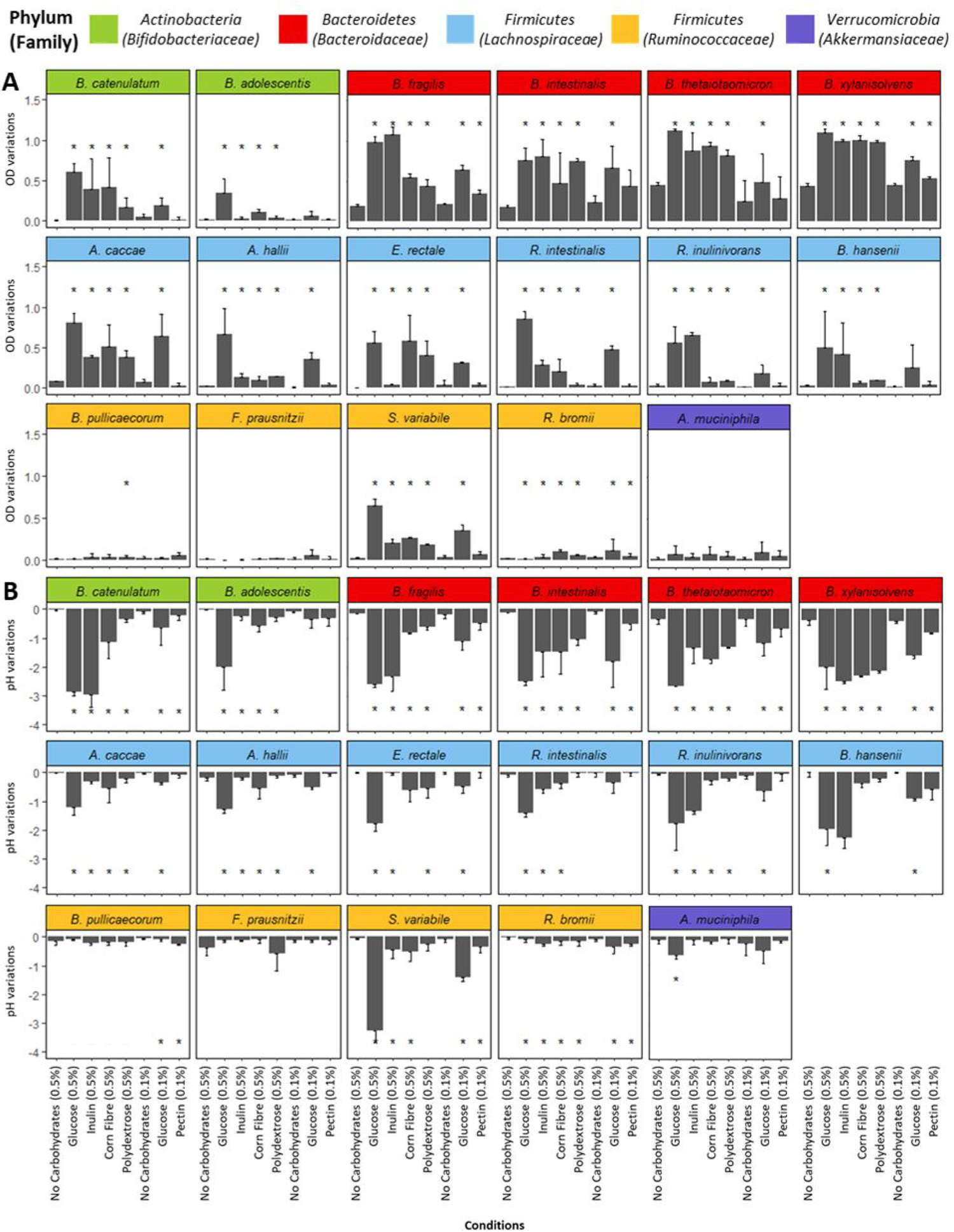
Functional activities of a selection of 17 commensal bacteria in the presence of carbohydrate substrates. Various carbohydrate sources were added into a low nutrient culture medium at concentrations of 0.5% (w/v for agave inulin, corn fiber, polydextrose) or 0.1% (w/v for citrus pectin), taking into account solubility and viscosity of dietary carbohydrates. Parallel controls included glucose at 0.5% or 0.1%, as well as a condition with no added carbohydrates. (A) Variations of OD at 600nm reflect bacterial growth after 24 h of culture. (B) Variations of pH reveal the medium acidification, as a consequence of the production of fermentation end-products, such as short-chain fatty acids, after 24 h of culture. OD and pH variations correspond to the difference between OD and pH values measured after 24 h of culture, in comparison to the value measured just after the inoculation of the bacteria. Comparisons were conducted within each bacterial species between different carbon sources and the non-supplemented medium using unpaired Mann-Whitney’s non-parametric tests. Mean values annotated with * are significantly different (p-value<0.0625) compared to the control condition of no carbohydrates.

### Effect of dietary carbohydrates on SCFA production

As major metabolites resulting from the breakdown of carbohydrates by gut bacteria, SCFA were quantified. Levels and types of SCFA varied greatly based on bacterial strains and the carbon sources (Fig. S1) (Text S1 for detailed SCFA analysis). A hierarchical classification method applied to SCFA production revealed four distinct fermentation profiles (Fig. 2). Cluster N, representing 64% of all monocultures, encompassed non-supplemented LNCM and LNCM-pectin conditions characterized by limited fermentation with low SCFA levels. This cluster mainly featured experiments involving *R. bromii*, *F. prausnitzii*, *B. pullicaecorum*, *B. adolescentis,* and *A. muciniphila*, demonstrating lower activities compared to the overall dataset. In contrast, the other clusters displayed intense fermentation activities with unique SCFA contents. Cluster G, representing 13% of all experiments, grouped bacterial strains that especially metabolise glucose, leading to significantly high butyrate concentrations averaging 8.9 mM ± 5.47. This cluster exclusively featured butyrate-producers from the *Firmicutes* phylum, particularly members of *Lachnospiraceae* family. This finding indicates that *E. rectale*, *A. caccae*, *R. intestinalis,* and *S. variabile* efficiently utilise glucose as a carbon source. Cluster A, representing 17% of all experiments, exclusively comprised *Bacteroidetes* strains, leading to significantly high propionate concentrations averaging 10.3 mM ± 5.20. Additionally, isovalerate production (mean of 2.1 mM ± 0.64) was associated with cluster A, highlighting the capacities of *Bacteroidetes* strains to breakdown peptides present in LNCM. Interestingly, this cluster (A) excluded the LNCM-glucose condition, supporting the hypothesis of an evolutive adaptation of *Bacteroidetes* species towards complex carbohydrate utilisation. Cluster G/I, representing 6% of all experiments, consisted of bacterial strains exhibiting high fermentation capacities in both LNCM-glucose and LNCM-inulin. This cluster regrouped *B. catenulatum*, *B. hansenii*, *B. fragilis,* and *B. xylanisolvens*, leading to significantly high acetate concentrations averaging 26.6 mM ± 7. In summary, the hierarchical clustering of SCFA profiles revealed four fermentation groups related to carbohydrate utilisation, partly reflecting the phylogenetic affiliation of commensal bacteria. Butyrate and propionate distinguished clusters G and A, closely associated with *Firmicutes* and *Bacteroidetes*, respectively. Acetate differentiated cluster G/I, which encompassed three bacterial phyla. Statistical analysis indicated that bacterial strains were the primary factor influencing carbohydrate utilisation, surpassing the impact of experimental conditions (Table S2).

**Figure 2:**
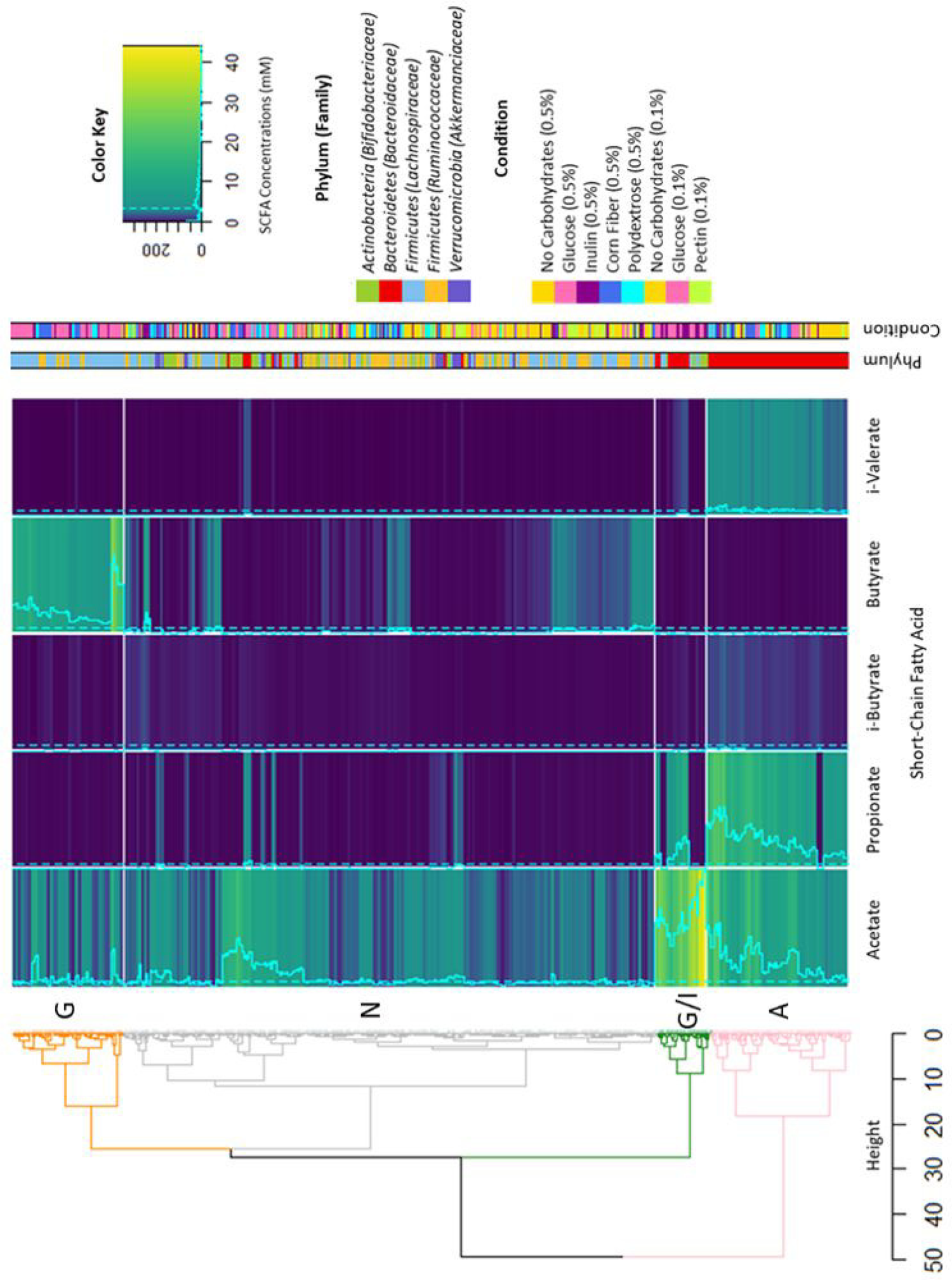
Classification of carbohydrate fermentation profiles of 17 commensal bacteria. In the dendrogram, all bacterial monocultures are grouped based on their fermentation profiles. We can observe four distinct clusters highlighting the functional capacities of commensal bacteria to ferment various carbohydrates. The blue shading represents short-chain fatty acids (SCFA) concentrations, expressed in mM. Dotted lines denote the overall mean of total SCFA, excluding valerate, i-caproate, and caproate due to their null concentrations. The straight lines indicate the SCFA values for each bacterial culture. The hierarchical classification identified four clusters: in grey, cluster N; in orange, cluster G; in pink, cluster A; in green, cluster G/I.

### Metabolomic profiles of six gut bacteria in the presence of dietary carbohydrates

The metabolic activities of six bacterial strains, representing *Bacteroidetes* and *Firmicutes* were investigated in the presence of various substrates using high-resolution liquid chromatography-mass spectrometry (HR LC-MS). The selected *Bacteroidetes* and *Firmicutes* strains, namely *B. xylanisolvens*, *B. thetaiotaomicron*, *B. intestinalis*, *S. variabile*, *R. intestinalis*, and *E. rectale*, were chosen for their capacity to grow in our experimental conditions (Fig.1). *B. xylanisolvens*, *B. thetaiotaomicron*, and *B. intestinalis* demonstrated robust growth on all carbon sources, while *S. variabile*, *R. intestinalis*, and *E. rectale* exhibited slight but significant growth with complex dietary carbohydrates. To evaluate their metabolic responses, we compared the metabolomic profiles obtained in non-supplemented LNCM, and those supplemented with simple (glucose) and complex (agave inulin, corn fiber, and citrus pectin) carbon sources against corresponding non-inoculated LCNM, considered as the reference conditions (Fig. 3A). First, sparse partial least squares-discriminant analyses (sPLS-DA) for each bacterial strains were computed based on the culture conditions. In the *Bacteroidetes* group, the metabolomic profiles clearly distinguished between inoculated *versus* non-inoculated media, even in non-supplemented LNCM (Fig. 3B and C, and D). However, in the presence of dietary carbohydrates, the metabolic patterns differed from those obtained in non-supplemented LNCM, indicating a shift in metabolism associated with complex sugars (Fig. 3B and C). Interestingly, the metabolic responses of *B. thetaiotaomicron* varied between conditions with glucose and complex dietary carbohydrates. While the separation of metabolomes in the presence of complex sugars was less pronounced for *B. intestinalis*, there were some distinctions, particularly with LNCM-inulin and LNCM-corn fiber (Fig. 3D). In the *Firmicutes* group, low metabolic activities were observed in non-supplemented LNCM clustering with non-inoculated conditions, and a metabolic shift occurred in the presence of glucose and complex sugars (Fig. 3E, F, and G). While the separation of metabolomes was less distinct than that observed for *Bacteroidetes* members, the presence of dietary carbohydrates induced a consistent metabolic shift across bacterial strains. The comparison of all conditions and bacterial strains confirmed the separation of metabolomes between *Bacteroidetes* and *Firmicutes* members (Fig. S2). Then, analysis focused on analytes with changed relative abundances in incubated LNCM versus non-inoculated LNCM, emphasizing metabolites with high-confidence annotation (Fig. 4, File S1) and those with predicted identities using public databases (Fig. S3, File S2) (Text S1 for detailed metabolite identification). Regardless of the condition, *Bacteroidetes* profiles consistently differed from *Firmicutes*, with *B. xylanisolvens* and *B. thetaiotaomicron* consistently clustering together (Fig. 4, Fig. S3). *B. intestinalis* exhibited the most distinctive metabolomic signature among *Bacteroidetes* members, while *E. rectale*, *R. intestinalis*, and *S. variabile* tended to cluster together under all conditions within the *Firmicutes* group (Fig. 4, Fig. S3). The key discriminating metabolite between the two phyla was succinate, with increased production by *Bacteroidetes* in the presence of glucose and complex dietary carbohydrates. Other *Bacteroidetes*–specific metabolites, such as hydroxyl-butyric acid, amino-isobutyric acid, and 2-hydroxy-hexadecanoic acid, increased with complex sugars. Some metabolites were specifically stimulated by dietary carbohydrates like sedoheptulose-7-phosphate, mainly induced by *Bacteroidetes* in LNCM-inulin, and O-phosphoryl-ethanolamine production, induced by *B. intestinalis* and *B. thetaiotaomicron* only in the presence of dietary carbohydrates. Notably, *B. xylanisolvens* induced the production of gamma-aminobutyric acid (GABA). For *Firmicutes* members, some metabolites increased with both glucose and complex dietary carbohydrates, including N-acetyl-L-cysteine, while lactic acid production was observed for all three strains in the presence of complex sugars. Strain-specific metabolites, such as malonic acid unique to *S. variabile* with glucose and inulin, and thymine or 4-methyl-2-oxovaleric acid unique to *E. rectale* with glucose, were identified. Nicotinic acid was consistently produced by *S. variabile* and *E. rectale* under all culture conditions. Several metabolites were modulated in bacterial strains from both phyla, including dehydroascorbic acid/cis-aconitic acid and D-Ribose 5-phosphate, which increased with complex sugars.

**Figure 3:**
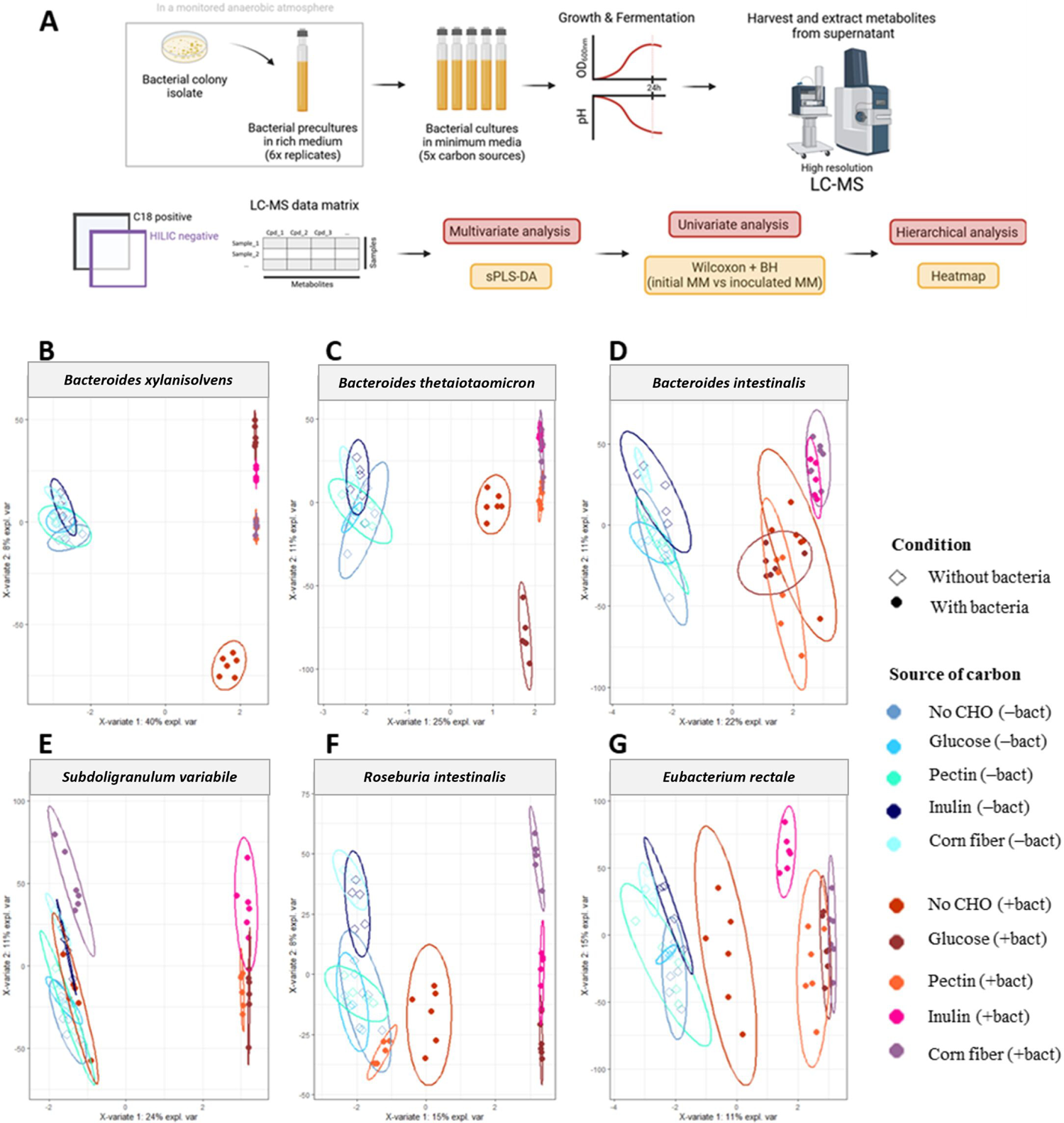
Metabolomic profiles of commensal bacteria according to culture condition and carbon sources. (A) Schematic of the experimental strategy and the untargeted metabolomic data analysis. The metabolomic study was conducted on supernatants of six commensal bacteria, belonging to two different bacterial phyla including *Bacteroidetes* (*B. xylanisolvens*, *B. thetaiotaomicron* and *B. intestinalis*) and *Firmicutes* (*S. variabile, R. intestinalis* and *E. rectale*). These bacterial species were cultivated in five low nutrient culture media (LNCM), each supplemented or not with different carbon sources. Each condition was performed in 6 replicates, in addition to 5 replicates of the initial non-inoculated LNCM. The LC-MS metabolomic approach consisted of two types of chromatographic conditions (C18 and HILIC), and two ionization conditions (both positive and negative modes) resulting in the detection of features in the HILIC column and negative ionization mode, and features in the C18 columns and positive ionization mode. Scatter plots of the first two sPLS-DA components were obtained for each of the bacterial species for all the culture condition (B) *B. xylanisolvens*; (C) *B. thetaiotaomicron*; (D) *B. intestinalis*; (E) *S. variabile*; (F) *R. intestinalis*; (G) *E. rectale*. All ellipses were drawn assuming a multivariate t-distribution with a confidence level of 0.95.

**Figure 4:**
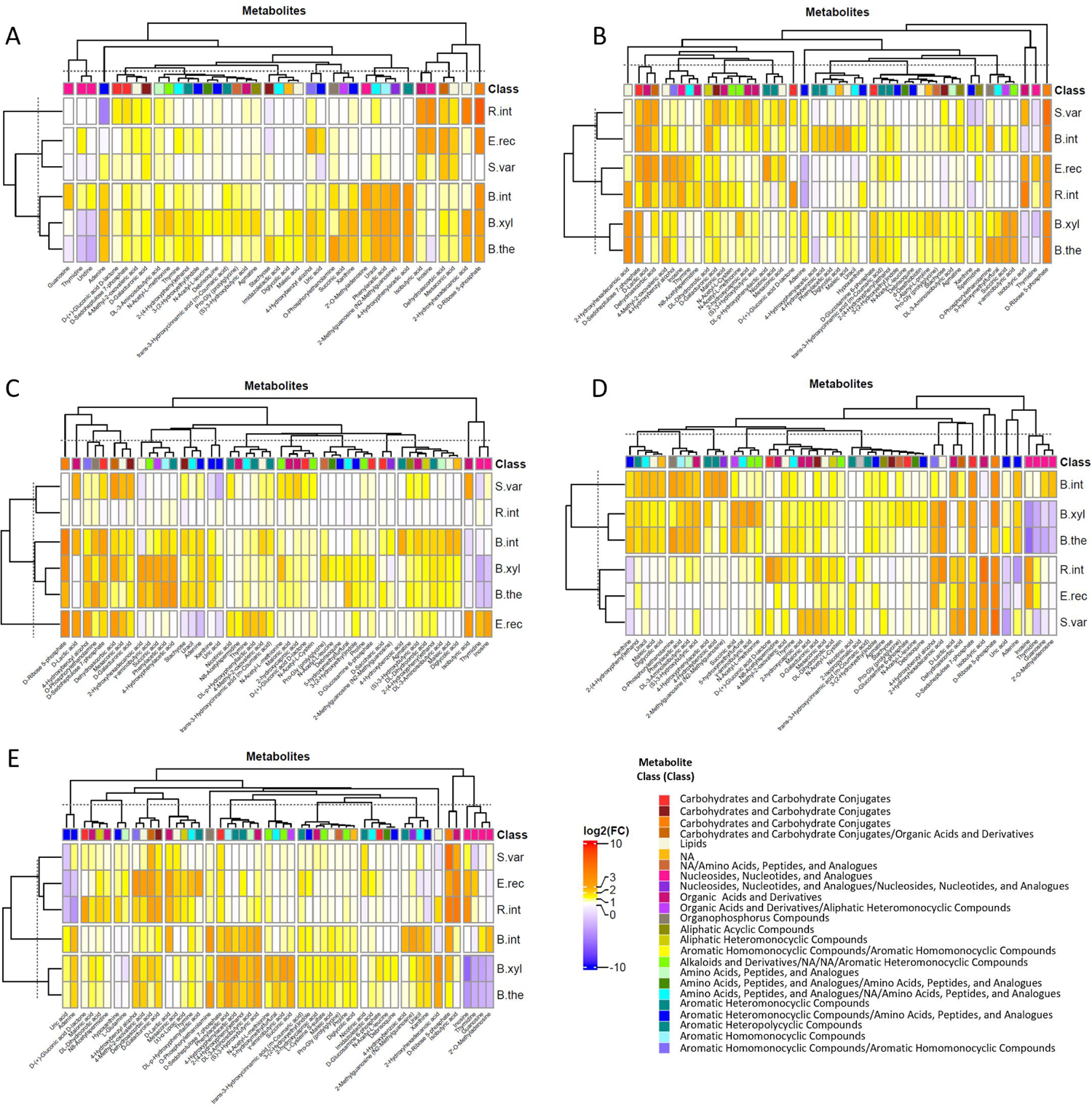
Heatmaps of differentially abundant metabolites after 24 h incubation of bacterial strains. **(A)** without any carbohydrate supplementation, (B) in the presence of 0.5% glucose, (C) 0.1% pectin, (D) 0.5% inulin, and (E) 0.5% corn fiber. The bacterial species were *B. xylanisolvens*, *B. thetaiotaomicron*, *B. intestinalis*, *S. variabile*, *R. intestinalis*, and *E. rectale* (named *B. xyla*, *B. theta*, *B. int*, *S. var*, *R. int*, *E. rect*, respectively). Metabolites that differ between medium inoculated with each of the bacterial species and the corresponding non-incubated medium were identified using univariate non-parametric test (Wilcoxon test, p<0.05). Metabolites plotted were significantly modulated in at least one condition with a fold change higher than 2. Only metabolites annotated with high level of confidence are shown.

### Molecular mechanisms underlying the interactions between the commensal bacteria and dietary carbohydrates

To elucidate the genetic responses to dietary carbohydrates, we conducted a molecular-scale investigation of the activities of four bacterial strains out of the six employed in the metabolomic analysis. This exploration occurred at the end of the exponential growth phase, focusing on three specific dietary carbohydrates (agave inulin, corn fiber, and citrus pectin) (Text S1 for detailed gene identification). The selection of bacteria for RNA-Seq analysis was guided by their substrate preferences to promote optimal growth. Specifically, we chose two *Bacteroidetes* strains, *B. thetaiotaomicron*, and *B. xylanisolvens*, whose growth showed robust stimulation in the presence of dietary carbohydrates. Additionally, we opted for two *Firmicutes* strains, *R. intestinalis* and *S. variabile*, which are known for their promising health-promoting properties and whose demonstrated growth responses to dietary carbohydrates. As a reference condition for saccharolytic mechanisms involving a simple carbon source, LNCM-glucose was used (Fig. 5A). First, differential expression analysis identified numerous genes significantly modified in LNCM-inulin and LNCM-corn fiber or LNCM-pectin (File S3). To capture the molecular responses of these bacterial strains, significant differentially expressed genes were affiliated with biological processes using gene ontology (GO) (Text S1). Enrichment tests of functional GO terms revealed that all bacterial transcriptomes were primarily associated with central metabolism related to nucleotides, amino acids, carbon, and energy conversion (Fig. S4). In *B. thetaiotaomicron*, the highest enrichment scores were associated with metabolic processes involving RNA in LNCM-inulin, DNA in LNCM-corn fiber, and organic acids (including pyruvate) in both conditions. *B. xylanisolvens* in LNCM-inulin showed enrichment in metabolic processes related to fatty acids, organic acids, and monosaccharides, which are likely involved in carbohydrate utilisation and energy conversion. In both LNCM-inulin and LNCM-corn fiber, this bacterium expressed genes associated with protein metabolism, including translation, metabolic processes of peptides, amides, and organonitrogen compounds. In LNCM-corn fiber, *B. xylanisolvens* highlighted genes involved in protein transport. Moreover, *R. intestinalis* exhibited enrichment in metabolic processes related to DNA and protein, including translation, metabolic processes of proteins, peptides, and amides. In LNCM-inulin, the molecular response was significantly associated with energy conversion and metabolic processes of fatty acids, monosaccharides, glucans, hexoses, and carbohydrate transport, suggesting carbohydrate utilisation. In LNCM-corn fiber, the GO profile was mainly affiliated with the metabolism of organic acids, alcohols, and vitamins, including thiamine and thiamine-containing compound biosynthesis. *S. variabile* displayed enrichment in energy conversion and metabolic processes involving DNA, lipids, organic acids, and carbohydrates in both LNCM. Particularly, in LNCM-pectin, the molecular response was significantly associated with aromatic compounds, heterocycles, and tetrapyrroles, likely related to cobalamin biosynthesis. Then, differentially expressed genes in response to various carbon sources among four selected bacteria revealed predicted CAZyme gene clusters (CGCs), each comprising at least one CAZyme, one transporter, one transcriptional regulator, and one signalling transduction protein [21]. Hierarchical clustering analysis of CGCs, combined across LNCM-inulin, LNCM-corn fiber, and LNCM-pectin, provided a comprehensive overview of biological functions related to carbohydrate utilisation (Fig. 5B). We focused on the top 5 CGCs with the most overexpressed genes for each LNCM. The transcriptomic response of *Bacteroidetes* strains included numerous genes annotated with functions encompassing various GH enzymes, transporters, and transcriptional factors. For instance, *B. xylanisolvens* overexpressed CGC2, which consisted of susC/susD-like, MFS transporter, and GH32 genes in LNCM-inulin, and displayed multiple upregulated genes, including susC/susD-like, GH13, GH29, GH92, GH97 in LNCM-corn fiber. *B. thetaiotaomicron* overexpressed CGC36, comprising susC/susD-like, MFS transporter, GH32, and GH18 genes in LNCM-inulin, and overexpressed CGC80, composed of susC/susD-like, GH13, GH97 genes, and CGC85, composed of susC/susD-like, GH18, GH92 genes in LNCM-corn fiber. In *Firmicutes* strains, the transcriptomic response showed numerous upregulated genes associated with transporters. *S. variabile* overexpressed CGC5, composed of GH3, GH32, GH2, LacI transcriptional factor, MFS, and PTS transporter genes, as well as CGC1, comprising GH77, LacI transcriptional factor, and ABC transporter genes in both LNCM-inulin and LNCM-pectin. *R. intestinalis* overexpressed genes encoding ABC transporters and ABC transporter permease in both LNCM. Notably, this bacterial strain exhibited upregulated genes involved in GH13, GH32, GH36, LacI transcriptional factor, and ABC transporters in LNCM-inulin.

**Figure 5:**
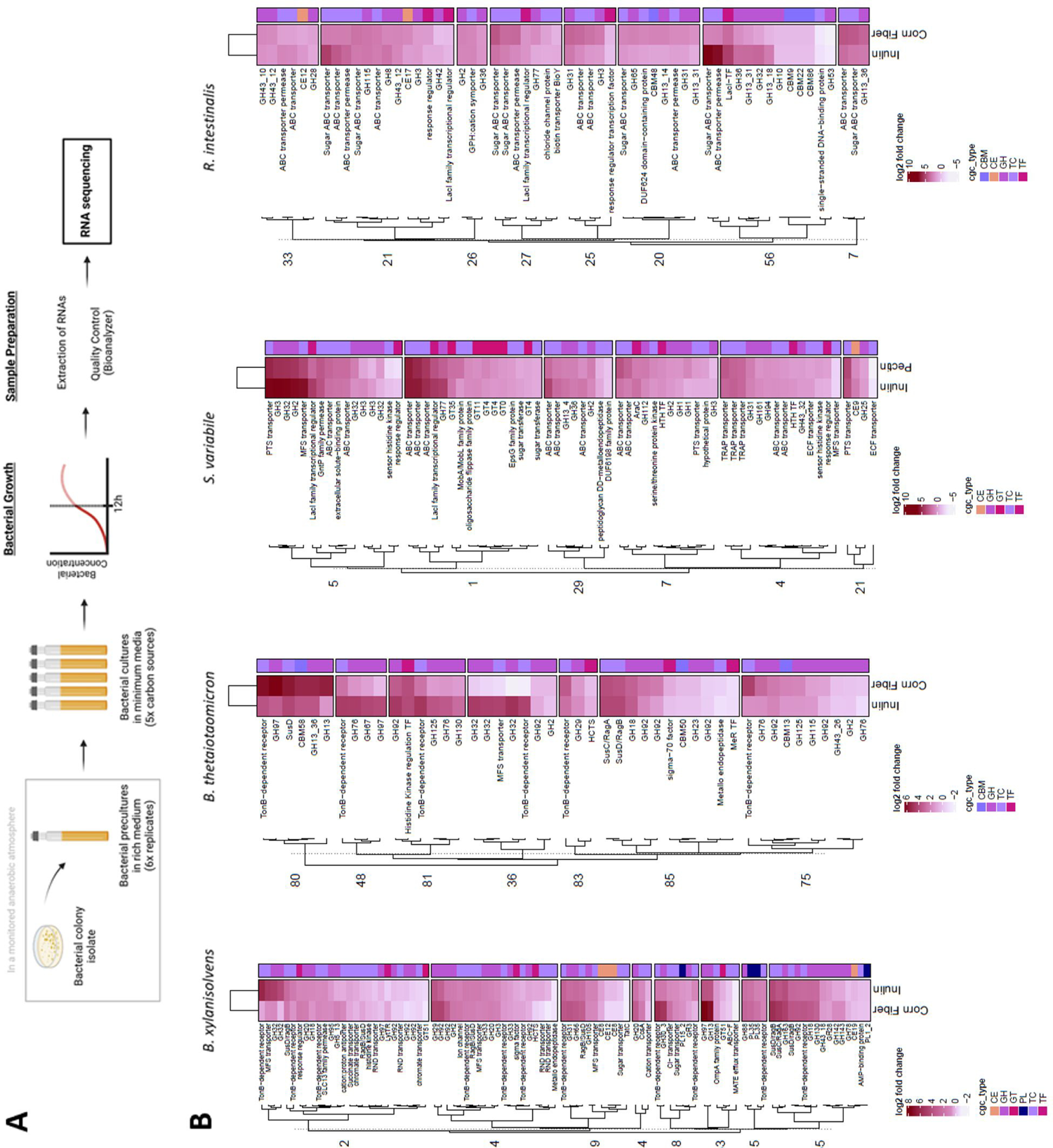
Transcriptomic profiles of four commensal bacteria in response to various carbon sources. (A) Schematic of the experimental strategy. The transcriptomic study was conducted on four commensal bacteria, belonging to two different bacterial phyla including *Bacteroidetes* (*B. thetaiotaomicron* and *B. xylanisolvens*) and *Firmicutes* (*R. intestinalis* and *S. variabile*). These bacterial species were cultivated in five low nutrient culture media (LNCM), each supplemented or not with different carbon sources. Each condition was performed in three replicates. (B) Hierarchical clustering heatmaps of expression changes for the most overexpressed genes regrouped in CAZyme gene clusters (CGCs) that were combined across the LNCM-inulin and LNCM-corn fiber or LNCM-pectin. GH: glycoside hydrolases; GT: glycosyl transferase; PL: polysaccharide lyases; CE: carbohydrate esterases; CBM: carbohydrate-binding module; TC: transporters; TF: transcription factors.

## DISCUSSION

The functional capacities of gut microbes to ferment dietary carbohydrates into metabolites within a dynamic ecosystem can shape the abundance and activities of various bacterial taxa, ultimately conferring beneficial effects on the host [22]. Nourishing the gut microbiota through prebiotics represents an important strategy to maintain and/or restore the bacterial community essential for gastrointestinal homeostasis [23]. Thoroughly characterising the capacities of commensal bacteria to metabolise dietary carbohydrates with prebiotic potential serves as an important basis for the rational development of nutritional and, more recently, clinical strategies [24]. Various approaches have provided insights into the impact of prebiotics on the gut microbiome, shedding light on carbohydrate utilisation mechanisms and physiological effects [25]. In this study, a panel of 17 commensal bacteria offered a lower-resolution view of the gut microbiota complexity. Our aim was to elucidate these bacterial capacities to metabolise four dietary carbohydrates with prebiotic potential. We anticipate that our research will serve as a benchmark for investigating carbohydrate-based prebiotic candidates, extending beyond the conventional focus on *Bifidobacterium* and *Lactobacillus* species, to target a broader spectrum of commensal bacteria.

### Growth and fermentation profiles revealed various degrees of carbohydrate utilisation

The carbohydrate metabolism of commensal bacteria was evaluated through growth and fermentation profiles. Bacterial taxa played a pivotal role in carbohydrate utilisation. *Bacteroidetes* strains demonstrated versatile carbohydrate utilization, particularly in acetate and propionate production (Fig. 2), attributed to their extensive CAZyme repertoires (Table S1) [15, 25, 26]. In contrast, *Actinobacteria, Firmicutes* and *Verrucomicrobia* strains displayed diverse profiles (Fig. 1), suggesting specialisation in a narrower range of complex dietary carbohydrates due to their limited CAZyme repertoires (Table S1) [27]. As emphasized in a previous study, the growth of commensal bacteria is likely dependent on both the bacterial strains and the polymerization degree of dietary carbohydrates [28, 29]. In our study, we opted to use consensus culture media, instead of employing optimized media for each strain. The objective was to identify the prebiotic potential of dietary substrates that enhance the growth of the bacterial strains under investigation. Thus, glucose served as the positive control for all strains, despite the fact that some strains, such as *B. adolescentis* and *B. catenulatum,* utilise glucose less than other substrate, like lactose [30]. Unfortunately, certain strains did not grow under any conditions (e.g. *B. pullicaecorum, F. prausnitzii, R. bromii* and *A. muciniphila*), possibly indicating suboptimal conditions with the chosen culture media.

### Bacterial responses to dietary carbohydrates were significant at the levels of metabolites

Beyond the SCFA production, little is known about the myriad of metabolites generated by carbohydrate-utilising bacteria, some of which may benefit the host [31]. Characterising these microbial molecules holds promise for clinical interventions and nutritional strategies [32]. Using untargeted HR LC-MS, we identified metabolic signatures that effectively discriminated *Firmicutes* and *Bacteroidetes* strains. Despite their close phylogenetic proximity in some instances, sPLS-DA analysis revealed distinct metabolomes for each bacterium, reflecting their adaptive responses to carbon source utilisation (Fig. 3, Fig. 4) [33]. Likely in line with their CAZyme repertoires, *Bacteroidetes* strains exhibited more active metabolisms, with certain analytes having potential health implications. For example, the production of succinate by *Bacteroidetes* strains is intriguing for host homeostasis and inflammation regulation [34], while the production of GABA, an inhibitory neurotransmitter, could impact anxiety and depression disorders in mammals [35]. In contrast, *Firmicutes* strains showed increased metabolite production in the presence of glucose and dietary carbohydrates, such as lactic acid [36]. Interestingly, nicotinic acid production by *S. variabile* and *E. rectale* irrespective of the culture condition, suggested a potential link to host nicotinamide adenine dinucleotide biosynthesis and a contribution to a symbiotic relationship in the gut microbiota [37]. Some metabolites were common to selected *Firmicutes* and *Bacteroidetes*, like dehydroascorbic acid/cis-aconitic acid and D-Ribose 5-phosphate, but annotating metabolites remains challenging [33]. Identifying unknown microbiota-derived metabolites and understanding their effects on human health remain significant research challenges [33]. Exploring these metabolic signatures could yield biomarkers characterising the impact of prebiotic dietary carbohydrates.

### Bacterial transcriptomes revealed the genetic traits involved in carbohydrate utilisation

Bacterial transcriptomes provided valuable insights into the metabolic functions involved in utilising diverse carbon sources. All four commensal bacterial strains activated genes related to central metabolic pathways. Notably, there were differential expressions of metabolic processes related to organic acids in all supplemented LNCM, indicating their essential role in bacterial activities. These organic acids, closely tied to carbohydrate metabolism, serve as carbon and energy sources, deriving from carbohydrate breakdown or other organic compounds. They contribute to ATP generation and the production of essential metabolites through respiration or fermentation. Moreover, the two *Firmicutes* strains exhibited differential gene expression involved in the metabolic processes related to B vitamins, including thiamine and cobalamin. Investigating the interplay between vitamin metabolism and fermentation activities could unveil novel mechanisms of action for carbohydrate-based prebiotics. The transcriptome profiles revealed distinct clusters of differentially expressed genes (Fig. 5). *Bacteroidetes* strains activated enzymes like CAZymes, susC-susD like transporters and transcriptional factors, resembling the archetypal structure of carbohydrate utilisation machinery [7, 16]. In contrast, *Firmicutes* strains prominently expressed genes encoding LacI transcriptional factors, CAZymes, and like ATP-binding cassette (ABC), phosphotransferase system (PTS), and major facilitator superfamily (MFS), suggesting the structural elements of Gram-positive PUL [7, 38]. The impacts of different carbon sources were discerned at the molecular scale for each bacterial strains. For instance, *Bacteroidetes* strains commonly activated CGCs encoding the most representative GH13, GH92 mannosidase, and GH97 glucoamylase in response to LNCM-corn fiber, indicating the selective degradation of this specific carbon source [26]. Similarly, CGCs containing GH32 were differentially expressed in all commensal bacteria in response to LNCM-inulin, underscoring the specificity of this enzyme in breaking down glycosidic bonds in inulin-type fructans [39]. To date, few studies have characterised the transcriptional activity of PUL-containing GH32 enzymes in LNCM-inulin, highlighting the selective utilisation of inulin by commensal bacteria. Our study provides molecular evidence of coordinated nutrient acquisition strategies in both *Bacteroidetes* and *Firmicutes* strains.

## CONCLUSION

In this study, not all carbohydrate substrates had an equal capacity to stimulate bacterial growth and fermentation activities. The observed variations in metabolic responses seemed to be shaped by the genetic characteristics of the commensal bacteria, with the phylogenetic affiliation of key intestinal microorganisms playing a pivotal role. In particular, *Bacteroidetes* strains demonstrated a broad capacity for carbohydrate degradation, in contrast to *Firmicutes* strains, which exhibited a more limited ability to utilise specific carbon sources. Comprehensive insights into bacterial responses were gained through metabolomic and transcriptomic analyses, revealing the underlying molecular mechanisms of carbohydrate metabolism at the strain level. While *in vitro* monoculture experiments offer valuable insights into prebiotic potential, heir translation to *in vivo* effects may be influenced by various factors that impact how prebiotics induce beneficial changes within the gut microbiome. Dietary interventions can represent a strategy to modulate the abundance and activity of beneficial bacteria in the gut [40]. Nevertheless, understanding carbohydrate utilisation capacities through a reductionist approach represents an initial step in the development of clinical interventions and nutritional strategies. The knowledge raised derived from our study establishes a foundation for potential applications aimed at enhancing gut health and overall well-being.

## MATERIALS & METHODS

### Dietary carbohydrates

This study included four dietary carbohydrates provided by General Mills. Agave-derived powdered long-chain inulin comprising fructose polymers (>90% dry weight) with minor amounts of fructose (<12% dry weight), glucose, and sucrose. Corn fiber consisted of a mixture of glucose polymers and insoluble non-digestible carbohydrates (>85% dry weight) derived from partially hydrolysed starch-made glucose syrup with minor monosaccharide content (<15% dry weight). Polydextrose comprising synthetic highly branched and randomly bonded polymers of glucose (>90% dry weight) with minor amounts of bound sorbitol and citric acid (<2% dry weight). Sourced from by-products of the juice and citrus-oil processing industries, citrus peel-derived pectin corresponds to low-ester pectin standardized with sugars.

### Bacterial strains, genome information and inoculum cultures

Bacterial strains were procured from *Deutsche Sammlung von Mikrooganismen und Zellkulturen* and the *American Type Culture Collection* (Table S1). Genomic data, either draft or complete, were retrieved from the National Center for Biotechnology Information database (https://www.ncbi.nlm.nih.gov) and subjected to quality assessment using QUAST and PROKKA [41, 42]. Selection of high-quality genomes met the following criteria: (1) only one genome per strain was selected; (2) among multiple genomes for the same strain, the genome with minimal number of contigs was selected; (3) if more than one genome with the maximum number of contigs exists, the genome with the maximal number of coding DNA sequence (CDS) was selected. Bacteria were isolated on supplemented BHI agar at 37°C within an anaerobic chamber containing 90% N_2_, 5% CO_2_, 5% H_2_ atmosphere (Coy Lab Products, USA) (Text S1). The identity of each isolate was systematically verified by 16S rRNA sequencing conducted by Eurofins. Colony PCR reactions were performed using a T100^TM^ Thermal Cycler (Biorad, Singapore) following manufacturer’s instructions with DreamTaq polymerase (Thermo Scientific, Lithuania), reverse primer [5’-ACGGCTACCTTGTTACGACTT-3’] and forward primer [5’-AGAGTTTGATCCTGGCTCAG-3’]. A single colony was cultured at 37°C for 24 h in supplemented BHI broth prior to be used as an inoculum culture diluted to an OD_600_ of 1 (Ultrapec 10 Cell Density Meter, Biochrom Ltd., UK). The counts of colony-forming units (CFU) were estimated to be in range of 1×10^7^ to 1×10^8^ CFU/mL.

### Culture conditions

A low nutrient culture medium (LNCM) was adapted from YCFA medium (Text S1) [43]. LNCM was supplemented with agave inulin, corn fiber or polydextrose at 0.5% (w/v) or citrus pectin at 0.1% (w/v) given the high viscosity of pectin solution at 0.5%. Carbon substrates were sterilised by 0.22 µm filtration. LNCM supplemented with glucose (at 0.5 or 0.1%) and with no additional carbohydrates were included as controls. The reconstituted culture media were allowed to equilibrate for at least 48h in the anaerobic chamber before being inoculated with 2% of the inoculum culture. We admit that nutrient carryover from pre-inoculations in rich media had only a marginal effect. Internal controls, devoid of biological material, were included. Metabolomic and transcriptomic experiments were conducted in Hungate tubes using single batches of LNCM adjusted to pH 6.8 and supplemented with or without carbon source, with a 20% reduction in the amounts of non-defined protein sources, such as Bacto^TM^ yeast extract and Bacto^TM^ tryptone.

### OD_600nm_ and pH measurements

Cultures were performed in duplicate in 2 mL 96-well plates, which were securely sealed with a sterile membrane (Thermo Scientific, USA) to prevent evaporation. Cultures were maintained under anaerobic conditions for 24 h at 37°C. Following homogenization of the samples, portions of each monoculture were transferred to 96-well plates (Corning, USA). Optical density (OD) at 600nm was measured both before (at time 0 - T0) and after (at time 24 – T24) bacterial growth using each non-inoculated LNCM as blanks (Tecan Infinite 200 Pro Plate Reader, Austria). Additionally, pH values were recorded before and after bacterial growth using a pH-meter 1140 (Mettler Toledo, Switzerland). To assess bacterial growth and the extent of carbohydrate utilization, we calculated OD and pH variations between T0 and T24 for each bacterial culture, where T0 is the value obtained just after the inoculation of the bacteria. All measurements were recorded as four biologically independent replicates. Data are reported as medians ± SD and analysed using R Studio v.4.0.0 [44]. For each bacterial strain, non-parametric Wilcoxon tests revealed pairwise differences between each carbon source and the non-supplemented medium after 24 h culture. Mean values annotated with * are significantly different (p≤0.0625) compared to the mean values in the absence of any carbohydrate supplementation.

### SCFA analysis

The heatmap was generated using the *heatmap.2* function from *gplots* v3.1.3. Plots were generated using *ggplot2* v3.4.1 [45]. SCFA content was conducted using gas chromatography (GC) equipped with flame ionization detection (FID) using the Agilent 7890 GC System (Text S1). All data were reported as medians ± SD in LNCM and analysed using R Studio v.4.0.0 [44]. For SCFA analysis, hierarchical clustering was performed using the *dendextend* v1.17.1 package. The clustering utilised the *hclust* function with the “ward.D2” agglomeration method and the *dist* function with the “Euclidean” method [46]. The hierarchical categories were described using the *FactoMineR* v2.8 package [47]. To visualize the results, the heatmap was generated with the *heatmap.2* function from the *gplots* v3.1.3 package. Plots were created using *ggplot2* v3.4.1 [44].

### Metabolomics

After 24 h incubation, samples were centrifuged at 12,000 g for 15 min at 4°C and supernatants were immediately stored frozen at −80°C until metabolomic analyses were performed using HR LC-MS (Text S1). Data matrices were analysed using R Studio v.4.0.0 [44]. Classification of metabolomic data was carried out using sPLS-DA, implemented in *mixOmics* v6.22.0 (Text S1) [48]. To identify significant features that distinguished between bacterial strains for a given carbon source, Wilcoxon tests were performed by comparing non-inoculated LNCM versus inoculated LNCM. Differentially abundant annotated metabolites were plotted on a heatmap after log2-transformation using *pheatmap* v1.0.12 [49].

### Transcriptomic sequencing and analysis

Bacterial cultures were harvested during the end of exponential growth phase, approximately 12 h after inoculation. Samples were centrifuged at 12,000 *g* for 15 min at 4°C and pellets were immediately frozen at −80°C until transcriptomic sequencing was performed using Illumina NextSeq500 (Text S1). Following transcriptomic data acquisition and bioinformatic analyses (Text S1), differential analyses were conducted using *edgeR* v3.38.4 [50]. Lists of differentially expressed genes were generated through likelihood ratio tests comparing LNCM-inulin and LNCM-corn fiber or LNCM-pectin, which relate to the utilisation of complex carbohydrates, versus the baseline LNCM-glucose. False discovery rate control was applied using Benjamini-Hochberg correction with a threshold of 0.05 [51]. To explore the relationships between genes and biological functions, gene ontology (GO) annotations encompassing biological process were employed using *ViSEAGO* v1.10 [52]. Functional enrichment analysis identified over-represented GO terms among differentially expressed genes at a significant level of 0.1 using the ‘classic’ algorithm with ‘fisher’ test. A minimal number of genes annotated to a GO term was settled at 5 using ‘genes_nodeSize’. For each LNCM, the mean of gene counts for the 3 replicates were considered. CGC were filtered to select at least one gene whose log2 fold change is greater than 5 for *Bacteroides* and greater than 2 for *Firmicutes*. Hierarchical classification was then performed for each LNCM using the agglomeration method ‘ward.D2’ and the ‘Euclidean’ method for distance calculation, as implemented in the *complexheatmap* v2.14.0 package.

## ACKNOWLEDGEMENTS

This work has benefited from the facilities and expertise of @BRIDGe (Université Paris-Saclay, INRAE, AgroParisTech, GABI, 78350, Jouy-en-Josas, France) and the high throughput sequencing core facility of I2BC (Centre de Recherche de Gif – http://www.i2bc.paris-saclay.fr/). We wish to thank the staff of the metabolomic platforms of MetaboHUB (Université Paris Saclay, INRAE, MetaboHUB, 91191 Gif-sur-Yvette, France) and Metatoul-AXIOM (INRAE, Toxalim, 31300 Saint-Martin du Touch, France). We wish to thank Anastats (https://www.anastats.fr/), in particular Sophie Dubois, for providing statistical consulting. We acknowledge the French PSPC (Projet Structurant Pour la Compétitivité; PSPC) RestorBiome project grant number DOS0099202/00 for their support. Figures were generated using BioRender (https://biorender.com).

## AUTHOR CONTRIBUTIONS

CBF, PB, SV and FP performed the experiments. FC and CCHO contributed to the metabolomic analyses. OR and CHA contributed to the transcriptomic analyses. CCHE and PL conceived and supervised the project. CBF wrote the first draft of the manuscript. CCHE, CCHA and PL corrected the manuscript. All the authors have read and accepted the final version of the manuscript.

## AVAILABILITY OF DATA

All summary data analysed in this study is included in supplementary files, and the datasets of OD, pH and SCFA in this study are available from the corresponding author upon reasonable request. The RNA-seq dataset used in this study can be found in NCBI, GSE237111.

